# ZGA: a flexible pipeline for read processing, de novo assembly and annotation of prokaryotic genomes

**DOI:** 10.1101/2021.04.27.441618

**Authors:** A.A. Korzhenkov

## Abstract

**Motivation:** Whole genome sequencing (WGS) became a routine method in modern days and may be applied to study a wide spectrum of scientific problems. Despite increasing availability of genome sequencing by itself, genome assembly and annotation could be a challenge for an inexperienced researcher.

**Results:** ZGA is a computational pipeline to assemble and annotate prokaryotic genomes. The pipeline supports several modern sequencing platforms and may be used for hybrid genome assembling. Resulting genome assembly is ready for deposition to an INSDC database or for further analysis.

**Availability:** ZGA was written in Python, the source code is freely available at https://github.com/laxeye/zga/. ZGA can be installed via Anaconda Cloud and Python Package Index.

**Supplementary information:** Supplementary data are available at *Bioinformatics* online.

## 1 Introduction

26 years have passed since the Haemophilus influenzae genome was sequenced in 1995 (Fleischmann, 1995) and high-throughput sequencing became trivial. To date (April 2021) NCBI RefSeq includes more than 200 000 genome assemblies of Bacteria and more than 1000 of Archaea. Genome Taxonomy Database (Release 06-RS202, 27th April 2021), which includes genomes of both cultivated and uncultivated prokaryotes, consists of 254 090 bacterial genomes representing 45 555 species of 12 037 genera and 4 316 archaeal genomes representing 2 339 species of 851 genera (Parks et al., 2020). Global prokaryotic diversity estimated to be one million OTUs (Louca et al., 2019) or 200 000 bacterial genera and 5 000 that of Archaea (Amann and Rosselló-Móra, 2016). Relying on these numbers, a large amount of novel genomic data will be obtained in future.

To date several instruments were developed for prokaryotic genome assembly and annotation (Supplementary Table S1), but most of them are abandoned. Old versions of assemblers integrated in the pipelines cannot handle data from modern sequencers (Nanopore, PacBio, BGI). Growth of existing databases, emergence of novel databases, development and modernisation of basic bioinformatic tools force use of updating annotation instruments. From pipelines specialized on full cycle processing of prokaryotic genomic data only few tools are still in active development: ASA3P (Schwengers et al., 2020), Bactopia (Petit and Read, 2020), nullarbor (Seeman et al., 2020), tormes (Quijada et al., 2019). These tools implement many features especially for comparative genomics while having their own limitations (Supplementary Table S1).

This work describes a novel computational pipeline for assembly and annotation of prokaryotic genomes using data from a wide variety of sequencing platforms.

## 2 Methods

Successful genome assembly and annotation requires several steps: read quality control, read pre-processing, genome assembling, genome quality assessment, genome annotation (Toshchakov et al., 2017). There are many tools which perform above mentioned operations so the following criteria were used for the selection:

- Computational efficiency
- Multithreading
- Accuracy and reproducibility
- Free use, availability of source code and permissive licensing
- Moderate system requirements not exceeding consumer PC
- Simplicity of integration
- Continuation of development
- Availability in software repositories.

Detailed description of pipeline steps and implemented software presented in Supplementary material 1.

### 2.1 Sequencing data

On 21 of April 2021 NCBI SRA database included 1,424,935 bacterial genome sequencing experiments conducted on different high-throughput platforms: Illumina (1,385,704), Pacific Biosciences SMRT (16,989), Oxford Nanopore (8,064), 454 Life Sciences (7,513), Ion Torrent (4,517), BGISEQ (1106), ABI SOLiD (724). This data shows that the developing pipeline should support popular modern platforms such as Illumina, Pacific Biosciences and Oxford Nanopore as well as emerging and prospective BGISEQ technology. Despite different technologies, sequencing data from Illumina and BGISEQ platforms are very similar in terms of read length, base quality distributions and overall sequencing quality (Kim et al., 2021). These two platforms use different library-prep adapters which should be trimmed during read processing. This issue was solved by addition of BGISEQ-specific adapter sequences to Illumina-oriented read processing workflow.

### 2.2 Genome assembly and quality assessment

To evaluate performance of ZGA pipeline several genomic datasets from NCBI database were selected for which both genome assembly (NCBI GenBank) and raw reads (NCBI SRA) were available (Supplementary table S4). Genomes with high and low GC-content were selected to show absence of limitations for the pipeline. Raw sequencing data had been processed with ZGA, Shovill, Bactopia and tormes. SPAdes (Bankevich et al., 2012) had been used for short read assemblies in all pipelines. nullarbor was unable to handle the data without a reference genome and had been excluded from comparison. QUAST (Gurevich et al., 2013) had been used to collect genome assembly metrics from available reference genomes and genome assemblies resulting from the pipeline.

### 2.3 Computational resources

The pipeline had been tested on several devices including laptop, PC and workstation. To emphasize the accessibility of the pipeline and it’s modest requirements I present results of pipeline run on a consumer laptop, specifications given in Supplementary Table S2. During the runs the memory limit was set to 8 GB to be close to modest volume of RAM in some laptops. The software versions are shown in Supplementary table S3.

## 3 Results

ZGA was written in Python and is compatible with interpreter versions from 3.6 and newer. The pipeline starts from long or short (paired or unpaired) sequencing reads and results in an annotated genome assembly. The pipeline may be stopped after any step to skip annotation, genome QC or genome assembly if only read-processing is desired. Processing of long reads is not implemented due to an internal correction algorithm of Flye. The pipeline workflow is shown on Figure 1.

**Fig. 1.**
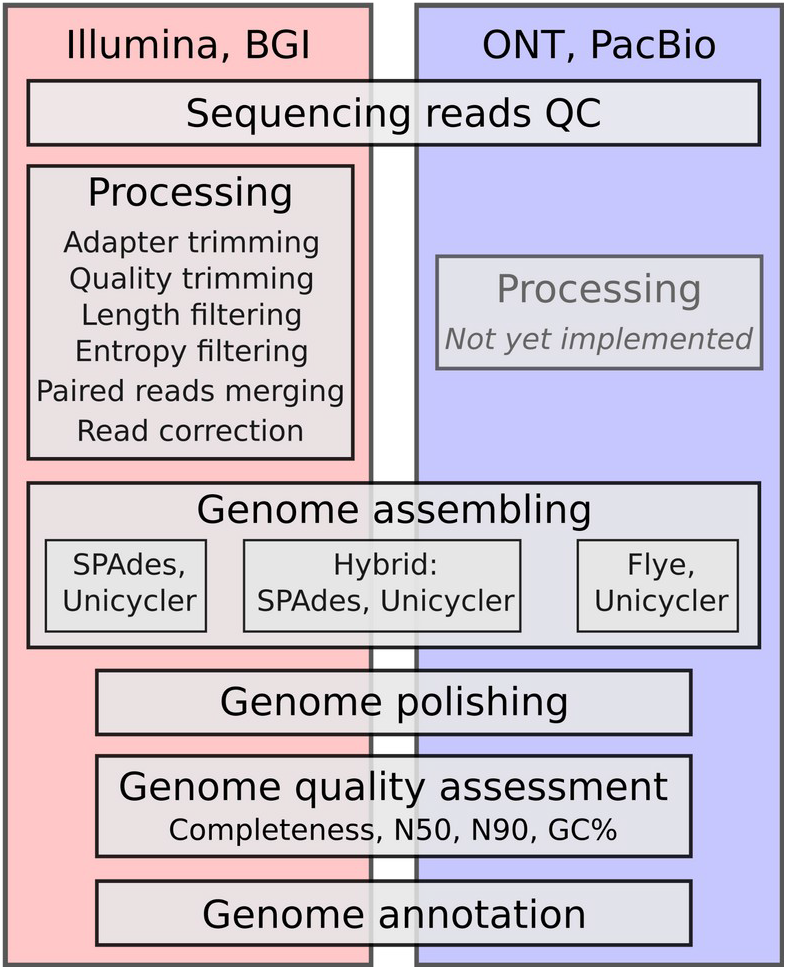
ZGA workflow representation.

All test datasets were successfully processed in a reasonable time, depending on the volume of sequencing data, except for tormes, which failed on BGI data (Supplementary Table S4). All short-read assemblies produced by ZGA show better or at least the same assembly quantitative metrics in comparison with assemblies deposited in the NCBI database. The pipelines were ranked by run time (less is better) and N50 metric (more is better) and contig count (less is better) of the resulting assemblies. ZGA and Shovill took the first place with 1.92 rank in average, Bactopia was third with 2.25, Tormes was fourth with 2.5.

## 4 Conclusion

ZGA pipeline successfully performs the key steps of prokaryotic genome assembly and annotation for data from different sequencing platforms. In less than an hour researchers may obtain an annotated genome assembly ready for further analysis or deposition in public databases. The source code structure allows tools replacement or addition to keep the pipeline using state-of-the-art tools and methods.

## Acknowledgements

The author thanks Katserov D. S. for manuscript proofreading and language correction and members of the Bioconda team for helping with recipe updating.

