## Supplementary Material 1 for "ZGA: a flexible pipeline for read processing, de novo assembly and annotation of prokaryotic genomes"

**Supplementary Material 1. Step of the genome assembly and annotation pipeline**

*1 Read quality control*

Read quality control (QC) performed with **fastp** (Chen et al., 2018) which supports both short and long sequencing reads and writes a report in HTML (user-friendly) and JSON (software-friendly) formats.

*2 Read processing*

Short read processing includes quality trimming, adapter trimming, filtering by length and complexity, merging overlapped reads. **BBTools** package (Bushnell, 2017) includes specialized tools for above mentioned actions: **BBDuk** for quality trimming, adapter removal and filtering by length, **BBMerge** for overlapping paired end reads merging and **filterbytile.sh** script for optional filtering of Illumina reads. Tadpole from BBtols package was included in the pipeline for kmer-based short read correction. Preliminary short read correction facilitates sequential read correction during genome assembly with SPAdes assembler. Correction and extension of short reads with **Tadpole** before SPAdes read-correction substantively lowers duration of the assembly process. In case of computation time or RAM usage constraints SPAdes default read-correction may be skipped altogether with reads being processed only through **Tadpole** before genome assembly.

Illumina mate-pair reads help with assembling larger scaffolds or even circular genomes. Mate-pair reads are processed with **NxTrim** (O'Connell et al., 2015) to filter out suspecting reads and trim adapter sequences. Long reads from ONT or PacBio platforms currently aren't processed due to the error correction stage in Flye assembler.

Genome size estimation may be useful for QC of assembled genome and is essential for genome assembling with Flye (Kolmogorov et al., 2019) and may be useful in QC. Genome size estimation in the pipeline is performed with **Mash** (Ondov et al., 2016).

*3 Assembly*

Three genome assemblers, such as SPAdes, Unicycler and Flye, were included in the pipeline for the assembly of prokaryotic genomes from both short sequencing reads (Illumina and BGISEQ platforms) and long sequencing reads (Pacific Biosciences and Oxford Nanopore Techonologies), and their combinations.

Three genome assemblers covering all possible variants of input files: short (Illumina, BGI) reads, long (PacBio, Nanopore) reads or their combination were selected: SPAdes, Unicycler and Flye.

**SPAdes** is a short-read genome assembler, which is able to use long reads for genome scaffolding (Bankevich et al., 2012). SPAdes includes a short read correction step, which may require high memory volume depending on genome size, genome GC-content, sequencing coverage and sequencing read quality. In case of insufficient memory volume on a user's computer this step may be skipped in the pipeline (not recommended) or substituted by Tadpole error correction. **Unicycler** assembler was intended for hybrid genome assembly, it uses SPAdes for short-read based assembly and **miniasm** for long-read based assembly (Wick et al., 2017). Unicycler performs extensive bridging that generally results in better assemblies than just using SPAdes at the cost of time. **Flye** assembler (Kolmogorov et al., 2019) works only with long reads, but the resulting assembly may be then polished using short reads in case they are provided. Flye needs to be provided with estimated genome size that may be specified by the user or calculated with Mash during the previous step of the pipeline.

*4 Assembly polishing*

Assembly polishing is a process of correcting bases, fixing mis-assemblies and filling gaps to improve quality of resulting genome assembly using sequencing reads. SPAdes includes MismatchCorrector, a module improving mismatch and short indel rates in the assembly which uses the **BWA** (Li and Durbin, 2009) and requires no additional polishing. Unicycler uses **Pilon** (Walker et al., 2014) for assembly improvement and also doesn't require external polishing. Flye assembler uses an internal tool to polish assembly with long reads. For a hybrid assembly with Flye there is a possibility to skip the internal stage of genome polishing with long reads and perform polishing with short reads using Racon (Vaser et al., 2017) which improves quality of the genome assembly.

*5 Assembly quality control*

PhiX is a sequencing control frequently added during Illumina sequencing and the study of Integrated Microbial Genomes database showed that more than 5% of genomes had PhiX contamination (Mukherjee et al., 2015). To detect PhiX contamination, a reference nucleotide sequence was aligned against genome assembly with BLASTn from NCBI BLAST+ (Camacho et al., 2009). Contigs which have alignment coverage of PhiX sequence more than 50% being removed from assembly using **Biopython** (Cock et al., 2009).

**CheckM** (Parks et al., 2015) is used to assess the quality of genome assembly: completeness, contamination and heterogeneity. Analysis is possible in two modes: taxonomy mode and lineage mode. In taxonomy mode the user provides rank and taxon name, otherwise the bacterial marker set is used. In lineage mode the marker set will be defined by CheckM, which requires more computational resources.

*6 Genome annotation*

To date several pipelines for prokaryotic genomes annotation are developed, but only two autonomous offline open-source tools are in active development: **Prokka** (Seeman, 2014) written in Perl and **DFAST** (Tanizawa et al., 2018) written in Python and supporting modern versions of the interpreter (ver. 3.6 and higher). To reduce dependencies count, decrease language heterogeneity and accounting for the announcement of Perl7 (https://www.perl.com/article/announcing-perl-7/) which may lead to broken dependencies of many Perl modules, including such used by Prokka, DFAST was implemented in ZGA pipeline.

*7 Output data*

After successful completion of the pipeline users obtain an annotated genome ready for submission to INSDC databases. Output directory also includes read quality statistics, processed reads, raw genome assembly and genome completeness and contamination report. For user convenience logging during the run of the pipeline was implemented.

*References*

Bankevich,A. et al. (2012) SPAdes: A New Genome Assembly Algorithm and Its Applications to Single-Cell Sequencing. Journal of Computational Biology, 19, 455–477.

Bushnell,B. et al. (2017) BBMerge – Accurate paired shotgun read merging via overlap. PLOS ONE, 12, e0185056.

Camacho,C. et al. (2009) BLAST+: architecture and applications. BMC Bioinformatics, 10, 421.

Chen,S. et al. (2018) fastp: an ultra-fast all-in-one FASTQ preprocessor. Bioinformatics, 34, i884–i890.

Cock,P.J.A. et al. (2009) Biopython: freely available Python tools for computational molecular biology and bioinformatics. Bioinformatics, 25, 1422–1423.

Kolmogorov,M. et al. (2019) Assembly of long, error-prone reads using repeat graphs. Nature Biotechnology, 37, 540–546.

Mukherjee,S. et al. (2015) Large-scale contamination of microbial isolate genomes by Illumina PhiX control. Standards in Genomic Sciences, 10, 18.

O’Connell,J. et al. (2015) NxTrim: optimized trimming of Illumina mate pair reads. Bioinformatics, 31, 2035–2037.

Ondov,B.D. et al. (2016) Mash: fast genome and metagenome distance estimation using MinHash. Genome Biology, 17, 132.

Parks,D.H. et al. (2015) CheckM: assessing the quality of microbial genomes recovered from isolates, single cells, and metagenomes. Genome Res., 25, 1043–1055.

Seemann,T. (2014) Prokka: rapid prokaryotic genome annotation. Bioinformatics, 30, 2068–2069.

Tanizawa,Y. et al. (2018) DFAST: a flexible prokaryotic genome annotation pipeline for faster genome publication. Bioinformatics, 34, 1037–1039.

Vaser,R. et al. (2017) Fast and accurate de novo genome assembly from long uncorrected reads. Genome Res., 27, 737–746.

Walker,B.J. et al. (2014) Pilon: An Integrated Tool for Comprehensive Microbial Variant Detection and Genome Assembly Improvement. PLOS ONE, 9, e112963.

Wick,R.R. et al. (2017) Unicycler: Resolving bacterial genome assemblies from short and long sequencing reads. PLOS Computational Biology, 13, e1005595.
