## Supplemental Table S1 for "ZGA: a flexible pipeline for read processing, de novo assembly and annotation of prokaryotic genomes"

**Supplementary Table S1. Computational pipelines for genome assembly and annotation.**

| | last update | long reads | hybrid assembly | mate-pair reads | read processing | genome size estimation | assembly | contig reordering | assembly polishing and correction | assembly QC | variant calling | genome annotation | taxonomical identification | MLST | pangenomics | report and visualization |
|---|---|---|---|---|---|---|---|---|---|---|---|---|---|---|---|---|
| **Tormes** | 2020 | - | - | - | + | - | Megahit, SPAdes | Mauve | - | - | - | Prokka | + | + | + | + |
| **ASA3P** | 2020 | + | + | - | + | - | SPAdes, HGAP, Unicycler | MeDuSa | - | + | + | Prokka | + | + | + | + |
| **Bactopia** | 2021 | Hybrid assembly only | + | - | + | + | Shovil | - | Pilon | + | + | Prokka | + | + | + | + |
| **Nullarbor** | 2020 | - | - | - | + | - | SKESA, SPAdes, Megahit, shovill, Velvet | - | - | - | + | Prokka | + | + | + | + |
| **iMetAMOS** | 2015 | PacBio only | - | + | + | - | 13 assemblers | - | - | 7 tools | - | Prokka | FCP, Kraken, Phylosift, PHMMER, PhymmBL | - | - | + |
| **MEGAnnotator** | 2015 | - | | | - | - | AbySS, MIRA, SPAdes | Mauve | - | + | - | | | - | - | |
| **MyPro** | 2014 | - | - | | + | | + | + | | | | | | | | |
| **A5-Miseq** | 2016 | - | - | | | | + | | | | | | | | | |
| **CloVR-Microbe** | 2017 | | | | | | | | | | | | | | | |
| **CG-Pipeline** | 2018 | | | - | | | Reference-based | | | | | + | | | | |
| **RAMPART** | 2016 | - | - | + | + | - | 7 assemblers, comparison and selection | - | Gap-closing and scaffolding | QUAST, CEGMA, KAT | - | | | | | - |
| **MAGpy** | 2019 | | | | | | | | | CheckM | - | + | Taxonomical annotation | - | - | + |
| **MetaSanity** | 2020 | | | | | | | | | CheckM | - | FuncSanity | GTDB-Tk, FastANI | - | - | + |
| **Shovill** | 2020 | - | - | - | + | + | SPAdes, SKESA, Megahit, Velvet | - | + | - | - | | | | | - |
| **ZGA** | 2021 | + | + | + | + | + | SPAdes, Unicycler, Flye | - | Polishing (Racon) | CheckM | - | DFAST | - | - | - | - |
