## Supplemental Table S2 for "ZGA: a flexible pipeline for read processing, de novo assembly and annotation of prokaryotic genomes"

**Supplementary table S2. Hardware specifications**

| Device | Description |
|---|---|
| CPU model | CPU: Intel(R) Core(TM) i5-10210U |
| Base CPU clock rate | 1.60 GHz |
| CPU cores/threads | 4/8 |
| CPU cache size | 6MB |
| RAM | 20 GB DDR4 |
| RAM speed | 2667MT/s |
| SDD | Hynix BC501 M.2 PCI-E NVMe 256Gb (HFM256GDJTNG-8310A) |
