## Supplemental Table S3 for "ZGA: a flexible pipeline for read processing, de novo assembly and annotation of prokaryotic genomes"

### Supplementary table S3. Software

| Software | Version |
|---|---|
| OS | Ubuntu 20.04.2 LTS |
| Linux kernel | 5.4.0.72 |
| Python | 3.7.6 |
| SPAdes | 3.14.0 |
| DFAST | 1.2.13 |
| conda | 4.10.1 |
| bbmap | 38.84 |
| blast | 2.9.0 |
| fasterq-dump | 2.10.9 |
| fastp | 0.20.1 |
| flye | 2.8.3 |
| unicycler | 0.4.8 |
| racon | 1.4.20 |
| mash | 2.0 |
| nxtrim | 0.4.3 |
| minimap2 | 2.17 |
| biopython | 1.76 |
| quast | 5.0.2 |
