## Supplemental Table S4 for "ZGA: a flexible pipeline for read processing, de novo assembly and annotation of prokaryotic genomes"

**Supplementary Table S4. Comparison of genome assembly and annotation pipelines**

| **NCBI SRA accession** | **SRR4114395** | **SRR5481474** | **SRR11434856** | **SRR11085890** | **SRR12342876, SRR12342877** |
|---|---|---|---|---|---|
| Sequencing techonology | Illumina | Illumina | BGI | BGI | Illumina and Nanopore |
| Data volume, GBp | 0,9 | 0,4 | 2.0 | 1.2 | 0.7 + 0.26 |
| GC content, % | 32.7 | 61.6 | 32.0 | 71.2 | 35.2 |
| NCBI assembly accession | GCA_001997145.1 | GCA_001999385.1 | GCA_012029995.1 | GCA_012163135.1 | GCA_013177495.2 |
| **NCBI deposited assembly** | | | | | |
| Assembly length, bp (> 500) | 2 765 545 | 2 436 796 | 2 459 081 | 5 305 674 | 5 695 148 |
| Contig count (>500 bp) | 32* | 76* | 43 | 52* | 3 |
| N50, bp | 137 411 | 64 590 | 167 171 | 685 653 | 5 182 254 |
| Largest contig, bp | 435 732 | 193 009 | 323 604 | 1 301 853 | 5 182 254 |
| **ZGA** | | | | | |
| Wall clock time, mm:ss | 22:16 | 9:05 | 73:06 | 76:08 | 21:59 |
| Assembly length, bp (> 500) | 2 768 272 | 2 445 737 | 2 456 738 | 5 299 891 | 5 693 997 |
| Contig count (>500 bp) | 27 | 70 | 43 | 27 | 4* |
| N50, bp | 330 244 | 129 002 | 167 171 | 949 601 | 5 176 137 |
| Largest contig, bp | 606 831 | 214 194 | 323 791 | 1 266 467 | 5 176 137 |
| **Shovill** | | | | | |
| Wall clock time, mm:ss | 10:13.54 | 10:06.04 | 8:28.31 | 29:41.19 | N/A |
| Assembly length, bp (> 500) | 2 824 957 | 2 441 188 | 2 462 020 | 5 302 635 | N/A |
| Contig count (>500 bp) | 580 | 70 | 36 | 27 | N/A |
| N50, bp | 8 979 | 101 499 | 167 239 | 799 517 | N/A |
| Largest contig, bp | 50 146 | 214 290 | 497 828 | 1 266 667 | N/A |
| **Bactopia** | | | | | |
| Wall clock time, mm:ss | 16:54.80 | 11:24.21 | 30:41.00 | 40:40.23 | 289:39 |
| Assembly length, bp (> 500) | 2 756 661 | 2 425 758 | 2 452 706 | 5 293 414 | 5 681 601 |
| Contig count (>500 bp) | 30 | 79 | 33 | 76 | 8 |
| N50, bp | 171 089 | 73 314 | 256 579 | 109 422 | 5 151 146 |
| Largest contig, bp | 524 158 | 213 652 | 497 500 | 365 589 | 5 151 146 |
| **tormes** | | | | | |
| Wall clock time, mm:ss | 27:47.55 | 6:18.66 | Failed due to prinseq error | Failed due to prinseq error | N/A |
| Assembly length, bp (> 500) | 2 767 983 | 1 952 399 | | | N/A |
| Contig count (>500 bp) | 29 | 1190 | | | N/A |

| | | | | | |
|---|---|---|---|---|---|
| N50, bp | 330 244 | 2 050 | | | N/A |
| Largest contig, bp | 606 447 | 12 129 | | | N/A |
